## Supplementary Information for "A dynamic attractor network model of memory formation, reinforcement and forgetting"

### Supplementary Information Text

#### Network activity until convergence

In order to study further the convergence of the network, we ran longer simulations (10 times longer than the ones shown in the main text) with a network of $N = 100$ units.

First, we considered the case of "Assembly evolution", with one assembly of 10 out of $N = 100$ neurons stimulated for 70000 times with $f = 1/(60\ a.u.)$. We observe that the assembly size increases until reaching a plateau level at the size of 69 neurons (Figure S1). Most of the evolution of the assembly (specifically, until almost 70% of the final size) takes place in the first 10% of the stimulation phase (see inset in Figure S1, top left), which corresponds to the stimulation duration analyzed in Figure 5 of the main text. Therefore, with $f = 1/(60\ a.u.)$, the size of the stimulated assembly stabilizes without further recruiting the remaining neurons of the network. However, an assembly can also recruit all the units in the network if the frequency of stimulation is high enough. This is the case of a single assembly of 10 out of N=100 neurons stimulated with $f = 1/(30\ a.u.)$ for 100000 times (Figure S2). Importantly, we observe that, at the time of the 100000$^{th}$ stimulation, : i) the network activity is still controlled by the external stimulation (Figure S2, left); ii) the network connectivity (Figure S2, right) holds meaningful information reflecting the stimulation history (i.e. one and only one assembly stimulated at relatively high frequency for relatively long time).

Since we showed that a single assembly of 10 out of N=100 neurons stimulated for 70000 times with $f = 1/(60\ a.u.)$ reaches a size of 69 neurons (Figure S1), it is reasonable to expect that two assemblies stimulated with $f = 1/(60\ a.u.)$ would take all the neurons of the network. We tested this, particularly to check if the condition of orthogonality between assemblies subject to uncorrelated stimuli would hold even in the extreme condition of having all the network being part of one or the other assembly. Our results show that two assemblies stimulated at $f = 1/(60\ a.u.)$ overall take the whole network. However, the two assemblies: i) reach similar sizes (Figure S3, bottom), in line with the fact of being stimulated at the same frequency; ii) remain well separated (Figure S3, top right), in line with the fact of being subject to uncorrelated stimuli.

Considering all this, these results further support that, in our model, the runaway dynamics of Hebbian learning [1] is successfully limited, allowing the network to deviate its activity from baseline, encode meaningful information through the change of assembly sizes and then go back to baseline after stimulation.

### Scalability of the model

In none of the simulations analyzed in Figure 11 of the main text there was overlap of neurons between different assemblies (Fig. S4).

### Stability mechanisms

The mechanisms adopted to stabilize the system (hard bounds of the synaptic weights; weight decay; synaptic normalization; divisive normalization) were all necessary to ensure the functional behavior of the system (Fig. S5).

### Comparison of assembly evolution in different experimental paradigms

In order to compare the evolution of assemblies stimulated with the same frequency but in different paradigms (namely: single-memory network; 2 patterns stimulated with the same frequency; 2 patterns stimulated with different frequencies), we compared the results shown in Fig. 6A, Fig. 7A and Fig. 8A of the main text. We can observe that the sizes of assemblies stimulated with the same frequency evolved overall similarly in the different paradigms (Fig, S6A). It can be noted, however, that, in case of 2 assemblies stimulated with $f = 1/(60\, a.u.)$, the assembly growth of each of the assemblies stimulated with $f = 1/(60\, a.u.)$ was slower compared to the case of 1 assembly stimulated with $f = 1/(60\, a.u.)$ and to the case of 2 assemblies stimulated with $f_1 = 1/(60\, a.u.)$ and $f_2 = 1/(120\, a.u.)$ (Fig, S6B, top; ANOVA and post-hoc t-tests at three different stages: t = 1000 a.u.; t = 200000 a.u. and t = 420000 a.u.). This result seems reasonable considering that, in the case of 2 patterns stimulated with $f = 1/(60\, a.u.)$, there were two patterns stimulated with a relatively high frequency, both recruiting neurons, instead of only 1 pattern or 1 pattern with relatively high frequency and one with relatively low frequency. Regarding the evolution of the assemblies stimulated with $f = 1/(120\, a.u.)$, we found a significant difference, at the end of the simulations, in the sizes of the assemblies stimulated with $f = 1/(120\, a.u.)$ in a single-memory network compared to the case of networks with 2 patterns (Fig, S6B, bottom; t-test). This result is in line with what found for the assemblies stimulated with $f = 1/(60\, a.u.)$.

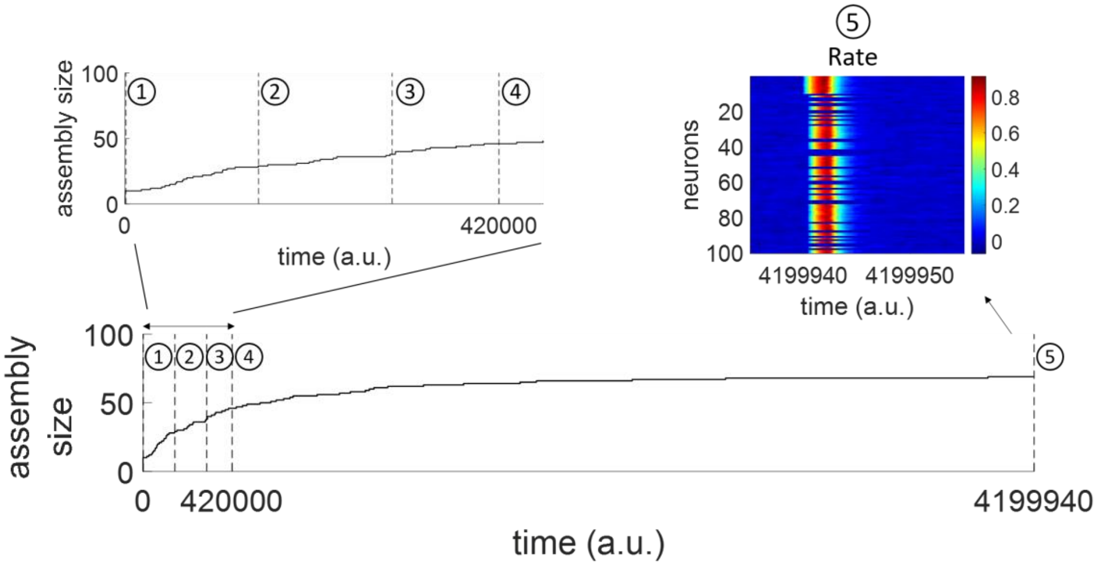


**Figure S1: Assembly evolution upon stimulus repetition until convergence.**

Starting at time 0 a.u., 10 neurons were stimulated for 70000 times with $\boldsymbol{f} = \mathbf{1}/(\mathbf{60}\ \boldsymbol{a.u.})$. Bottom**:** Number of assembly neurons over time. Inset, top left: zoom with enlarged time scale. Top, right: firing rate for all neurons at the time of the 70000$^{th}$ stimulation (the 10 directly stimulated neurons are on top).

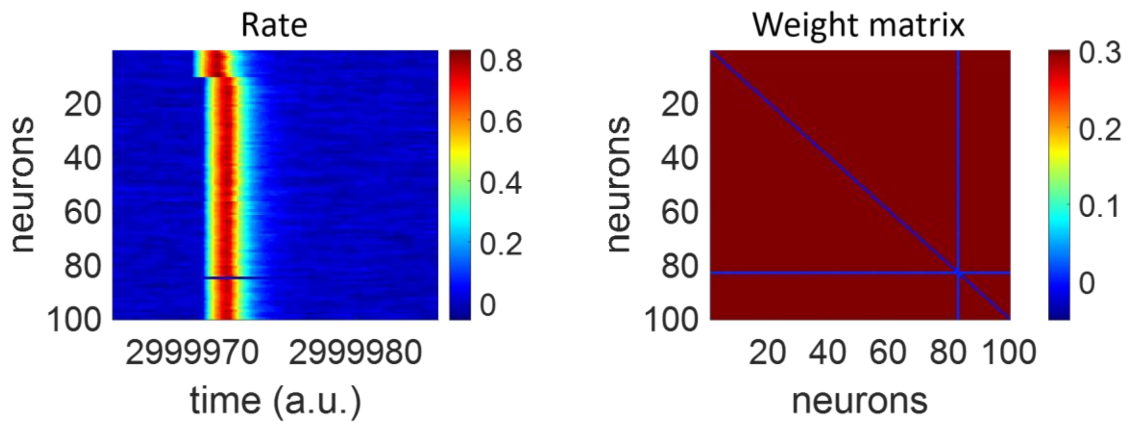


**Figure S2: Firing rate for all neurons and weight matrix at the time of the 100000$^{th}$ stimulation.**

Starting at time 0 a.u., 10 neurons were stimulated for 100000 times with $\boldsymbol{f} = \mathbf{1}/(\mathbf{30}\ \boldsymbol{a.u.})$. The firing rate plot and the weight matrix show that almost all the network had been recruited into one assembly after 100000 stimulations.

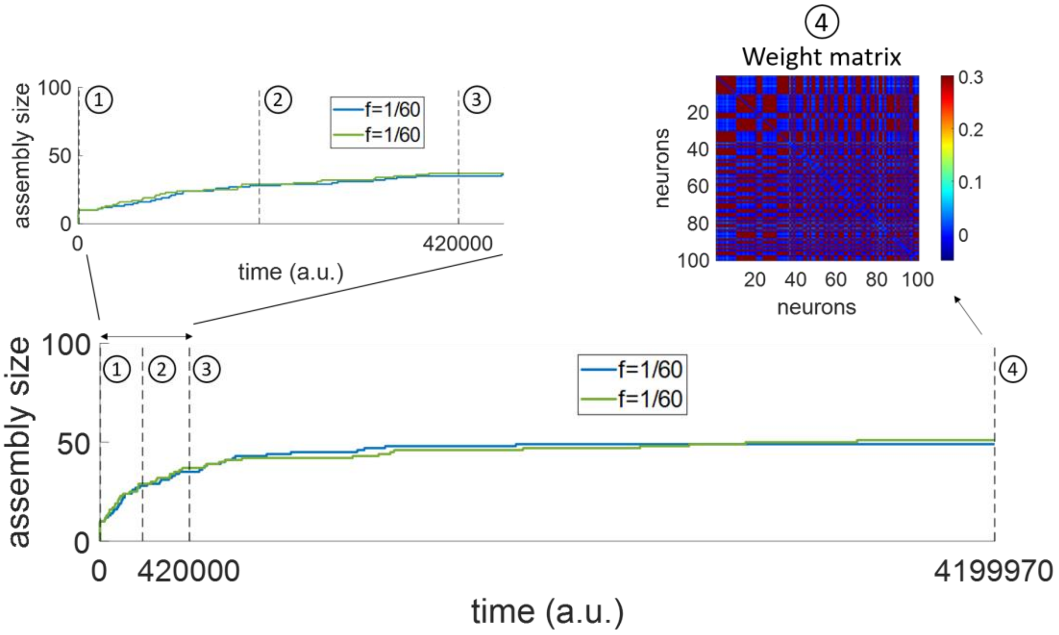


**Figure S3: Assembly evolution with two concurrent patterns stimulated at the same frequency until convergence.**

Starting at time 0 a.u., two non-overlapping populations of 10 neurons each were stimulated at different times with $f = 1/(60\ a.u.)$ for 70000 times. Bottom: number of neurons per assembly over time. Inset, top left: zoom with enlarged time scale. Top, right: weight matrix at the end of the stimulation paradigm.

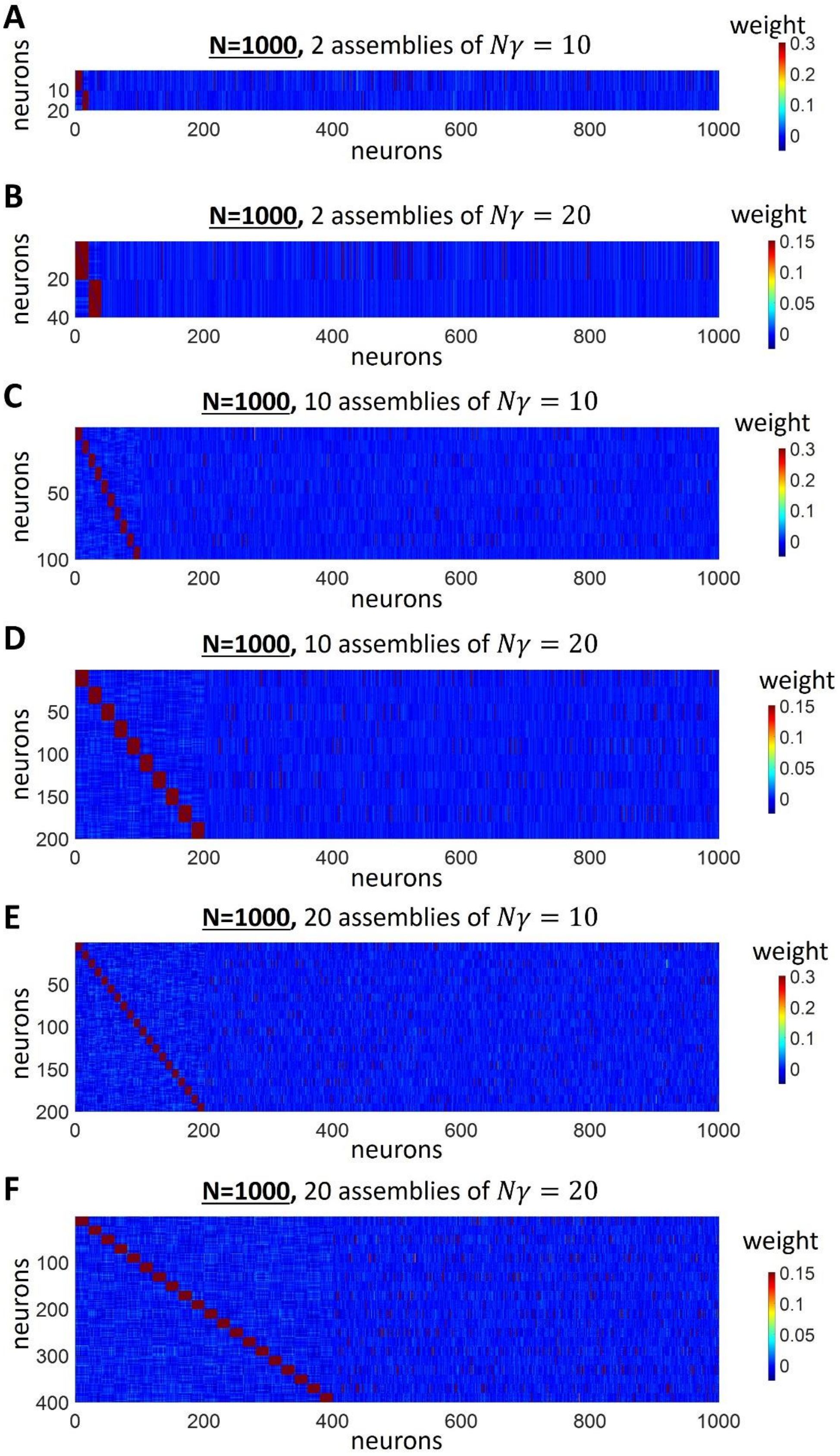
A
N=1000, 2 assemblies of $N\gamma = 10$
weight
neurons
B
N=1000, 2 assemblies of $N\gamma = 20$
C
N=1000, 10 assemblies of $N\gamma = 10$
D
N=1000, 10 assemblies of $N\gamma = 20$
E
N=1000, 20 assemblies of $N\gamma = 10$
F
N=1000, 20 assemblies of $N\gamma = 20$

**Figure S4: Connections among stimulated assemblies within networks of N=1000 with different combinations of number of assemblies and number of stimulated neurons.**

A-B-C-D-E-F) Connectivity, at t=350000 a.u., among stimulated assemblies within networks of 1000 neurons, in case of different number of assemblies and number of stimulated neurons, namely: A) 2 assemblies of 10 stimulated neurons ($N \cdot \gamma = 10$); B) 2 assemblies of 20 stimulated neurons; C) 10 assemblies of 10 stimulated neurons; D) 10 assemblies of 20 stimulated neurons; E) 20 assemblies of 10 stimulated neurons; F) 20 assemblies of 20 stimulated neurons. In each simulation, two stimulation frequencies ($f_1 = 1/(600\, a.u.)$ and $f_2 = 1/(1200\, a.u.)$) were used, with half of the assemblies stimulated with $f_1$ and half of the assemblies stimulated with $f_2$. For better visualization, only connections in one direction are shown (however, reciprocal connections have the same value). Connections among the non-stimulated neurons are not shown. In none of the simulations we observed the formation of overlaps between different assemblies.

(Parameters: $\beta = 0.000125$.)

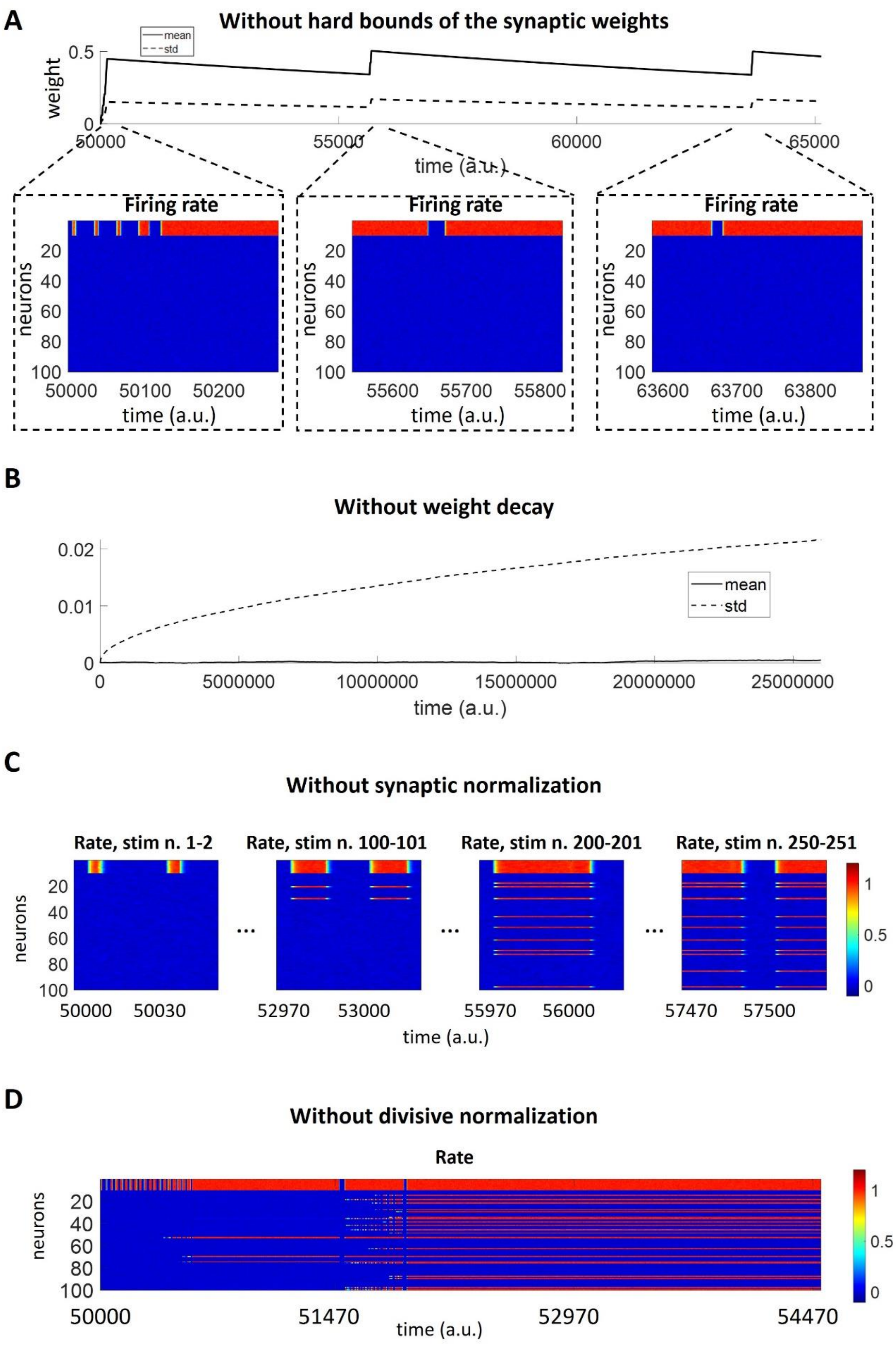
A
Without hard bounds of the synaptic weights
weight
mean
std
0.5
0
50000
55000
60000
65000
time (a.u.)
Firing rate
neurons
20
40
60
80
100
50000
50100
50200
55600
55700
55800
63600
63700
63800
time (a.u.)
B
Without weight decay
0.02
0.01
0
0
5000000
10000000
15000000
20000000
25000000
time (a.u.)
C
Without synaptic normalization
Rate, stim n. 1-2
Rate, stim n. 100-101
Rate, stim n. 200-201
Rate, stim n. 250-251
neurons
50000
50030
52970
53000
55970
56000
57470
57500
time (a.u.)
1
0.5
0
D
Without divisive normalization
Rate
neurons
50000
51470
52970
54470
time (a.u.)

**Figure S5: Model behaviour following selective removal of its stability mechanisms.**

A) Model without hard bounds of the synaptic weights. A specific population of 10 neurons was stimulated repeatedly at a repetition frequency $f = \frac{1}{30\ a.u.}$. The mean weight of the network increased without limit until the stimulated neurons would not return to baseline activation after the end of the external stimulation. The weight successively decreased, due to forgetting and due to the fact that sustained co-activation for longer than $T_{LR}$ (i.e. length of the window adopted for calculating the learning rule's running average; see Table 1 in the main text) does not result in learning (see Eq. 8 in the main text).

B) Model without weight decay. The mean and standard deviation of all network connections in absence of any external stimulation are displayed in the case of model without forgetting. Without forgetting, the standard deviation kept increasing, even if the other stability mechanisms were implemented.

(Parameters: $\beta = 0$).

C) Model without synaptic normalization: A specific population of 10 neurons was stimulated repeatedly at a repetition frequency $f = \frac{1}{30\ a.u.}$. The firing rates for all network neurons at different times are displayed. In absence of synaptic normalization, the formation of new connections with new recruited neurons lead to more prolonged assembly activation, eventually leading to uncontrolled network activity.

(Parameters: $\alpha_w = 0$; $\alpha_r = 2$).

D) Model without divisive normalization. A specific population of 10 neurons was stimulated repeatedly at a repetition frequency $f = \frac{1}{30\ a.u.}$. The firing rates for all network neurons are displayed, showing that, in absence of divisive normalization, the formation of new connections with new recruited neurons lead to uncontrolled network activity. (Parameters: $\alpha_w = 2$; $\alpha_r = 0$).

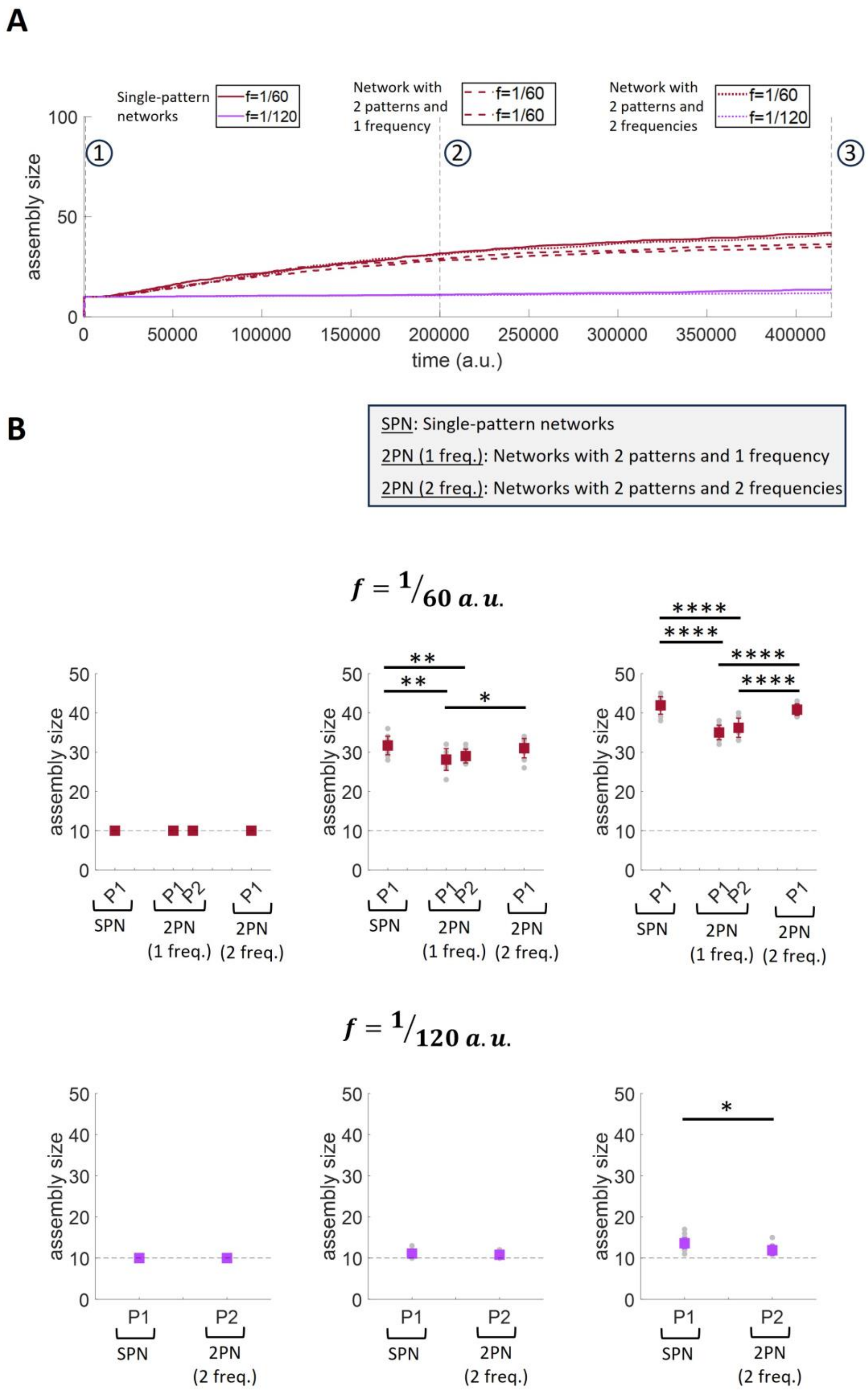
A
Single-pattern networks
f=1/60
f=1/120
Network with 2 patterns and 1 frequency
f=1/60
f=1/60
Network with 2 patterns and 2 frequencies
f=1/60
f=1/120
assembly size
time (a.u.)
B
SPN: Single-pattern networks
2PN (1 freq.): Networks with 2 patterns and 1 frequency
2PN (2 freq.): Networks with 2 patterns and 2 frequencies
f = 1/60 a.u.
f = 1/120 a.u.
P1
P2
SPN
2PN (1 freq.)
2PN (2 freq.)

**Figure S6: Comparison of assembly evolution for patterns stimulated with the same frequency in different experimental paradigms.**

A) Number of neurons per assembly over time in case of “Single-pattern networks”, “Network with 2 patterns and 1 frequency”, “Networks with 2 patterns and 2 frequencies”.

B) Number of neurons per assembly at different times for each of two stimulation frequencies ($f = \frac{1}{60\ a.u.}; = \frac{1}{120\ a.u.}$) in three different experimental paradigms. It should be noted that the assemblies’ names “P1” and “P2” correspond, for each paradigm, to the ones used in the figures of the main text (Fig. 6A, Fig. 7A and Fig. 8A).
